# TGFβ regulation of perilacunar/canalicular remodeling is sexually dimorphic

**DOI:** 10.1101/737395

**Authors:** Neha S. Dole, Cristal S. Yee, Courtney M. Mazur, Claire Acevedo, Tamara Alliston

## Abstract

Bone fragility is the product of defects in bone mass and bone quality, both of which show sex-specific differences. Despite this, the cellular and molecular mechanisms underpinning the sexually dimorphic control of bone quality remain unclear, limiting our ability to effectively prevent fractures, especially in postmenopausal osteoporosis. Recently, using male mice, we found that systemic or osteocyte-intrinsic inhibition of TGFβ signaling, achieved using the 9.6-kb DMP1 promoter-driven Cre recombinase (TβRII^ocy−/−^ mice), suppresses osteocyte perilacunar/canalicular remodeling (PLR) and compromises bone quality. Since systemic TGFβ inhibition more robustly increases bone mass in female than male mice, we postulated that sex-specific differences in bone quality could likewise result, in part, from dimorphic regulation of PLR by TGFβ. Moreover, since lactation induces PLR, we examined the effect of TGFβ inhibition on the female skeleton during lactation. In contrast to males, female mice that possess an osteocyte-intrinsic defect in TGFβ signaling were protected from TGFβ-dependent defects in PLR and bone quality. The expression of requisite PLR enzymes, the lacuno-canalicular network, and the flexural strength of female TβRII^ocy−/−^ bone was intact. With lactation, however, bone loss, and induction in PLR and osteocytic parathyroid hormone type I receptor (PTHR1) expression, were suppressed in TβRII^ocy−/−^ bone, relative to wild-type. Indeed, differential control of PTHR1 expression, by TGFβ and other factors, may contribute to dimorphism in PLR regulation in male and female TβRII^ocy−/−^ mice. These findings provide key insights into the sex-based differences in osteocyte PLR that underlie bone quality and highlight TGFβ signaling as a crucial regulator of lactation-induced PLR.

## 2. Introduction

Evolution selects for skeletal adaptations that improve survival and reproductive fitness. A prime example is the ability of female mammals to release stored calcium from bone to support lactation. In spite of a sharp drop in bone mass during lactation, the female skeleton mechanically adapts to accommodate these metabolic demands^(1–5)^. Changes in bone quality, including trabecular microarchitecture and cortical bone geometry, compensate for the loss of bone mass. Other aspects of bone quality, including the material properties of bone extracellular matrix (ECM), show sexual dimorphism^(6,7)^. Through mechanisms that remain unclear, the protective advantage of such skeletal adaptations for reproductive females is lost later in life. Women over 50 have a 4-fold higher rate of osteoporosis relative to men of the same age^(8,9)^. Furthermore, about half of osteoporotic fractures cannot be explained by clinically low bone mass^(10,11)^. Therefore, understanding the sexually dimorphic regulation of bone quality will elucidate mechanisms of bone fragility in post-menopausal osteoporosis, which affects over 200 million people worldwide^(12,13)^.

Skeletal variation in men and women arises in part from endocrine differences in hormones such as androgens, estrogens, and inhibin^(14)^. These hormones act in concert with paracrine growth factors to control bone cell function^(15,16)^. Osteoblasts and osteoclasts also exhibit sexual dimorphism at the cell-intrinsic level^(17–20)^. Osteocytes respond to high levels of PTH/PTHrP during lactation by inducing perilacunar/canalicular remodeling (PLR)^(21,22)^. In PLR, osteocytes resorb, and later replace, bone matrix surrounding their intricate lacuno-canalicular network^(21,23–25)^. Originally termed osteocyte osteolysis, PLR supports the metabolic demand for calcium during lactation through coordinated induction of several key genes, including matrix metalloproteinases (Mmps; namely Mmp2, Mmp13, and Mmp14), cathepsin K (Ctsk), carbonic anhydrase 2, and tartrate-resistant acid phosphatase (Acp5/TRAP) to mediate resorption^(23,24,26–28)^. After weaning, osteocytes replace the bone lost during lactation by inducing soluble cytosolic phosphatase (Phospho1) and dentin matrix protein (Dmp1)^(29–31)^. Until recently, PLR was primarily considered a response to metabolic stress or disease. Work from our group and others reveals PLR to be crucial in homeostasis as well, where it regulates bone quality through TGFβ signaling^(32)^. However, the extent to which the sexually dimorphic control of PLR extends dimorphism in bone quality remains unknown.

TGFβ is a key driver of bone mass and quality that dictates activities of all of the cells of the bone remodeling unit. Indeed, dysregulated TGFβ signaling plays a causal role in several skeletal diseases, including Camurati Engelman disease and osteogenesis imperfecta^(33,34)^. This key cytokine, which is largely produced by osteoblasts and osteocytes, is embedded within the bone ECM in a latent form^(35–37)^. Upon osteoclastic resorption, TGFβ is released and activated to promote recruitment and commitment of osteoblast progenitors to the resorption site, while inhibiting their terminal differentiation. TGFβ also promotes osteoclast formation and survival and couples bone resorption with formation^(38)^. In osteocytes, we found that TGFβ signaling is indispensable for PLR^(32,39)^. Using pharmacologic TGFβ antagonists and a novel mouse model, we demonstrated that osteocyte-intrinsic ablation of TGFβ signaling is sufficient to suppress PLR and cause bone fragility due to severely compromised bone quality^(32)^. Although female mice accrue more bone mass in response to pharmacologic TGFβ antagonists than males^(36)^, the sexually dimorphic regulation of PLR or bone quality by TGFβ has not been evaluated.

Therefore, we used mice with an osteocyte-specific deletion of TGFβ receptor II (TβRII) to investigate the sexually dimorphic regulation of PLR in mice at homeostasis and during the metabolic stress of lactation. Our findings reveal a dual role for TGFβ in the control of PLR in female bone. While TGFβ is required for lactation-induced PLR, at homeostasis, female bone is insensitive to loss of osteocytic TGFβ signaling. Here we elucidate a cellular mechanism that contributes to the dimorphic control of PLR in males and females, a mechanism that protects bone quality in reproductive age females.

## 3. Material and Methods

### 3.1 Mice

The generation of viable osteocyte specific TβRII^ocy−/−^ mice was described previously^31^. In brief, homozygous TGFβ type II receptor (TβRII)-floxed mice were bred with the hemizygous 9.6-kb DMP1-Cre mice^39,40^. Half of the mice resulting from the cross were DMP1-Cre^+/−^; TβRII^fl/fl^ (named TβRII^ocy−/−^ mice) and half were DMP1-Cre^−/−^; TβRII^fl/fl^ littermate controls (named Wild-type (WT) mice). For genotyping PCR, 50 ng of genomic DNA isolated from tail biopsies were used as a template. The presence of a Dmp1-Cre coding sequence was confirmed using forward 5’-TTG CCT TTC TCT CCA CAG GT-3’, reverse 5’-CAT GTC CAT CAG GTT CTT GC-3’ primers; the presence of a TβRII floxed allele was confirmed using the following primers, forward 5’-TAT GGA CTG GCT GCT TTT GTA TTC-3’, reverse, 5’-TGG GGA TAG AGG TAG AAA GAC ATA-3’. All the transgenic mouse lines were backcrossed for 8 generations into a congenic C57BL/6 background. In all the experiments 15-week old male and female TβRII^ocy−/−^ and WT mice were used.

For lactation studies, TβRII^ocy−/−^ mice and WT control littermates were time-mated at 10 weeks of age. Females that did not become pregnant after 1 week were removed from the study to minimize age-related differences in the bone phenotype. Following delivery, for each lactating mother the litter size was adjusted to 8 pups in order to normalize for breast milk requirements. Mice were sacrificed after 7 days of lactation. Baseline measurements were performed on age-matched virgin cohorts. Like males, female TβRII^ocy−/−^ and WT mice at 15-weeks of age were utilized in the study.

Mice, housed in a pathogen-free facility at 22°C with a 12-hour light/dark cycle, were supplied with standard irradiated mouse chow and water ad libitum. All studies were conducted with the approval of the Institutional Animal Care and Use Committee of the University of California San Francisco.

### 3.2 Micro-computed tomography (micro-CT)

For skeletal phenotyping, 15-week-old male and female mouse bones were harvested by cleaning off the surrounding soft tissue. Cleaned bones were subsequently fixed in 10% neutral buffered formalin and stored in 70% ethanol. A 2 mm region of the femoral metaphysis and a 1 mm region of the femoral mid-diaphysis were scanned using a Scanco µCT50 specimen scanner to assess trabecular and cortical parameters, respectively. Scanning was performed with an X-ray potential of 55 kVp and current of 109 µA and at a resolution of 10 µm. After scanning, scan projections were reconstructed to generate cross-sectional images using a cone-beam reconstruction algorithm. Density equivalent values are measured by calibrating the scanner to a hydroxyapatite phantom provided by the manufacturer. The trabecular bone compartment (300 µm proximal to epiphyseal plate) was segregated using manual delineation of the endosteal surface and morphometry was characterized by measuring the bone volume fraction (BV/TV), trabecular thickness (Tb. Th.), trabecular number (Tb. N.), and trabecular spacing (Tb. Sp.). Cortical morphometry was quantified within a 500 µm mid-diaphyseal span (50 serial sections) centered at the mid-point between proximal and distal epiphyses. Cross-sectional measurements included cortical bone area (Ct. Ar.), thickness (Ct. Th) and mineralization (Ct. Min). Table 1 shows standard parameters for n≥8 mice per group^41^.

**Table 1:** Micro CT data demonstrating trabecular and cortical bone parameters of male and virgin and lactating female WT and TβRII^ocy−/−^ mice at 15-weeks of age. *p<0.05 significantly different from the male WT virgin group ^a^ p<0.05 significantly different from the WT virgin group ^b^ p<0.05 significantly different from the WT lactation group

|  | Male |  | Female |  | Lactation |  |
| --- | --- | --- | --- | --- | --- | --- |
| | WT | TβRII<sup>ocy-/-</sup> | WT | TβRII<sup>ocy-/-</sup> | WT | TβRII<sup>ocy-/-</sup> |
| **Distal Femur** |  |  |  |  |  |  |
| **Tb. BV/TV (%)** | 0.18±0.04 | 0.30±0.07* | 0.08±0.04 | 0.1±0.05 | 0.04±0.01<sup>a</sup> | 0.08±0.02<sup>a,b</sup> |
| **Tb.N (1/mm)** | 5.30±1.08 | 6.11 ±0.87 | 2.44±0.72 | 2.68±1.13 | 1.76±0.45 | 2.89±0.84 |
| **Tb. Th (μm)** | 0.03±0.003 | 0.05±0.01* | 0.033±0.01 | 0.035±0.01 | 0.022±0.002<sup>a</sup> | 0.0271±0.00<sup>a,b</sup> |
| **Tb. Sp (mm)** | 0.16±0.05 | 0.12±0.03* | 0.40±0.12 | 0.42±0.22 | 0.59±0.2<sup>a</sup> | 0.35±0.15<sup>b</sup> |
| **SMI** | 1.71±0.36 | 0.52±0.62* | 2.83±0.43 | 2.74±0.66 | 3.01±0.27 | 2.48±0.38 |
| **Tb. Min (mg HA/cm<sup>3</sup>)** | 1065.48±18.9 | 1098.31±16.3* | 1103.85±30.2 | 1096.5±23.6 | 1010.5±27.0<sup>a</sup> | 1035.25±40.9<sup>c</sup> |
| **Midshaft Femur** |  |  |  |  |  |  |
| **Ct. BA/TA (%)** | 0.28±0.03 | 0.30±0.03 | 0.28±0.02 | 0.29±0.02 | 0.21±0.03<sup>a</sup> | 0.24±0.02<sup>a,b</sup> |
| **Ct. Th** | 0.20±0.01 | 0.22±0.02* | 0.19±0.01 | 0.20±0.01 | 0.14±0.006<sup>a</sup> | 0.16±0.007 |
| **Ct. SMI** | 0.16±0.41 | 2.82±1.42* | 0.28±0.14 | 0.12±0.23 | 0.22±0.03 | 0.10±0.18 |
| **Ct. Min (mg HA/cm<sup>3</sup>)** | 1385.81±14.4 | 1337.81±18.7* | 1362.60±46.6 | 1332.5±66.4 | 1330.3±29.1 | 1319.16±36.5 |

### 3.3 Quantitative RT-PCR analysis

As previously described, RNA was extracted and purified from bones using the miRNeasy mini kit (Qiagen, Valencia, CA)^31^. Briefly, soft tissue was cleaned off of bones (humeri) and marrow was flushed before RNA extraction. Following quantification, cDNA was synthesized from total RNA using the iScript cDNA synthesis kit (BioRad). Real-time quantitative PCR (qPCR) was performed using specific primers (iQ SYBR Green Supermix, BioRad) and Taqman probes (Applied Biosystems Taqman Assays, Thermo Fisher Scientific) following manufacturer’s protocol. Results are shown as relative gene expression, using expression of the 18s ribosomal RNA as an internal reference gene. Sequences of the primers have been provided in a table in the *Supplemental data* (*Table S1*).

### 3.4 Histology

Femurs from the TβRII^ocy−/−^ and WT male and female virgin and lactating mice (15 weeks old) were dissected, fixed in 10% neutral buffered formalin and decalcified in 10% di- and tetra-sodium EDTA for 28 days. Bone were then processed for paraffin embedding and sectioning using a microtome. 7μm thick sections were used for immunohistochemistry and Ploton silver stain.

For immunohistochemistry, sections were incubated with rabbit polyclonal anti-MMP13 antibody (1:100; Abcam ab39012); rabbit polyclonal monoclonal anti-MMP14 antibody (1:100; Abcam ab53712); rabbit polyclonal anti-Cathepsin K (CTSK) antibody (1:75; Abcam ab19027), or mouse monoclonal anti-TβRII antibody (1:500; Abcam ab186838) and mouse monoclonal anti-PTHR1 antibody (1:25, NBP1-40067: Novus Biologicals). Subsequently, sections were incubated with HRP-conjugated secondary antibody and developed with a DAB-substrate chromogen system (Innovex Universal Animal IHC kit). Cells expressing the protein of interest are stained brown. Corresponding mouse or rabbit negative control sera was used to verify the specificity of primary antibodies.

Ploton silver staining was performed to visualize the osteocyte lacuno-canalicular network in bones and the osteocyte network area was determined and normalized to total bone area, as previously described^31,42^. Osteocyte number was evaluated using hematoxylin and eosin staining. Images were acquired using a Nikon Eclipse E800 bright-field microscope. The number of osteocytes that stained positive in a given bone area were assessed using ImageJ (Rasband, W.S., ImageJ, U.S. National Institutes of Health, Bethesda http://imagej.nih.gov/ij/). The quantitative average represents data from 4-5 high-powered fields for each bone. Sections were evaluated for one femur from each of n≥3 mice per group, as indicated in the figure legends.

### 3.5 Synchrotron radiation Micro-Tomography (SRµCT)

SRµCT studies were used to assess the degree of mineralization of the bone as well as the volume and degree of anisotropy of the lacunae. The mid-diaphysis of 15-week-old virgin and lactating female mouse femora were scanned with a 20 keV x-ray energy, a 300 ms exposure time and a 5x magnifying lens for a spatial resolution of 1.3 µm (n = 4 bones/group). Imaging was performed at the Advanced Light Source on Beamline 8.3.2 at the Lawrence Berkeley National Laboratory by obtaining two-dimensional radiographs as the specimens were rotated from 0° to 180°. The radiographs were reconstructed in 3D Fourier-filtered back projection using Octopus software (Octopus v8; IIC UGent, Zwijnaarde). The attenuation coefficient or gray values for each voxel is divided by the mass attenuation coefficient of bone (4.001 cm^2^/g at 20 keV) to directly calculate bone mineral density. Avizo (VSG, Visualization Sciences Group) visualization and analysis software was used to filter, segment, and quantify the lacunar features.

### 3.6 Flexural strength tests

Whole bone flexural strength of femora was determined by macromechanical testing of intact, hydrated femurs isolated from TβRII^ocy−/−^ and WT mice. 15-week old males and female virgin and lactating mice of each genotype were used for macromechanical testing as previously described^31^. Intact femurs were loaded in three-point bending on a 6 mm span with the posterior side in tension using a Bose Electroforce 3200 test frame. For all tests, load-displacement data were recorded at 10 Hz. Bone cross sections were subsequently imaged in a scanning electron microscope (Zeiss Sigma 500 VP FE-SEM) for calculation of moment of inertia. Bending modulus was calculated from the linear portion of the stress-strain curve using standard beam theory equations, and yield stress was calculated using the 0.2% offset method.

### 3.7 Serum Analysis

Blood was isolated from male and virgin and lactating female mice using a cardiac puncture protocol and processed for serum. Serum calcium and phosphorus were measured with colorimetric assays (Abcam). PTH in serum was analyzed by Enzyme-linked Immunosorbent Assay (ELISA) (Quidel Corporation, Catalog no. 60-2305).

### 3.8 Statistical Analysis

Sample size was determined based on a power calculation that provides an 80% chance of detecting a significant difference (p<0.05). Technical replicates and biological replicates (n) used for all experiments are described in the figure legends. Where technical replicates are used, data are expressed as mean ± S.E.M. Otherwise, data are reported as mean ± S.D. Prism 5.0 (GraphPad Software, Inc., San Diego, CA, USA) was used for statistical analysis. Comparisons between two groups were evaluated by an unpaired two-tailed Student’s t test. For comparing between more than two groups, we used one-way ANOVA followed by Tukey’s test for multiple comparisons.

## 4. Results

### 4.1. Loss of osteocyte-intrinsic TGFβ signaling impacts bone mass and quality in male but not female mice

Recent studies from our group implicate osteocyte-intrinsic TGFβ signaling to be key for controlling perilacunar/canalicular remodeling (PLR) and bone quality, however it is unclear if this regulatory function of TGFβ is subject to sex-specific differences. Indeed, we previously observed sexual dimorphism in the effect of TGFβ inhibition on bone mass and osteoblast and osteoclast numbers^(36)^. To address this question, we used a previously developed mouse model for osteocyte-intrinsic inhibition of TGFβ signaling to assess the bone phenotype in both sexes. Using the 9.6-kb Dmp1-Cre transgene, the floxed TβRII allele was preferentially excised from odontoblasts and osteocytes of TβRII^ocy−/−^ mice^(32,41,42)^. The efficiency of this mouse model was confirmed by observing reduction in TβRII mRNA and protein levels in the long bone osteocytes of TβRII^ocy−/−^ male and female mice (Figure 1A-D, Fig S1A, Fig 3D).

**Figure 1.**
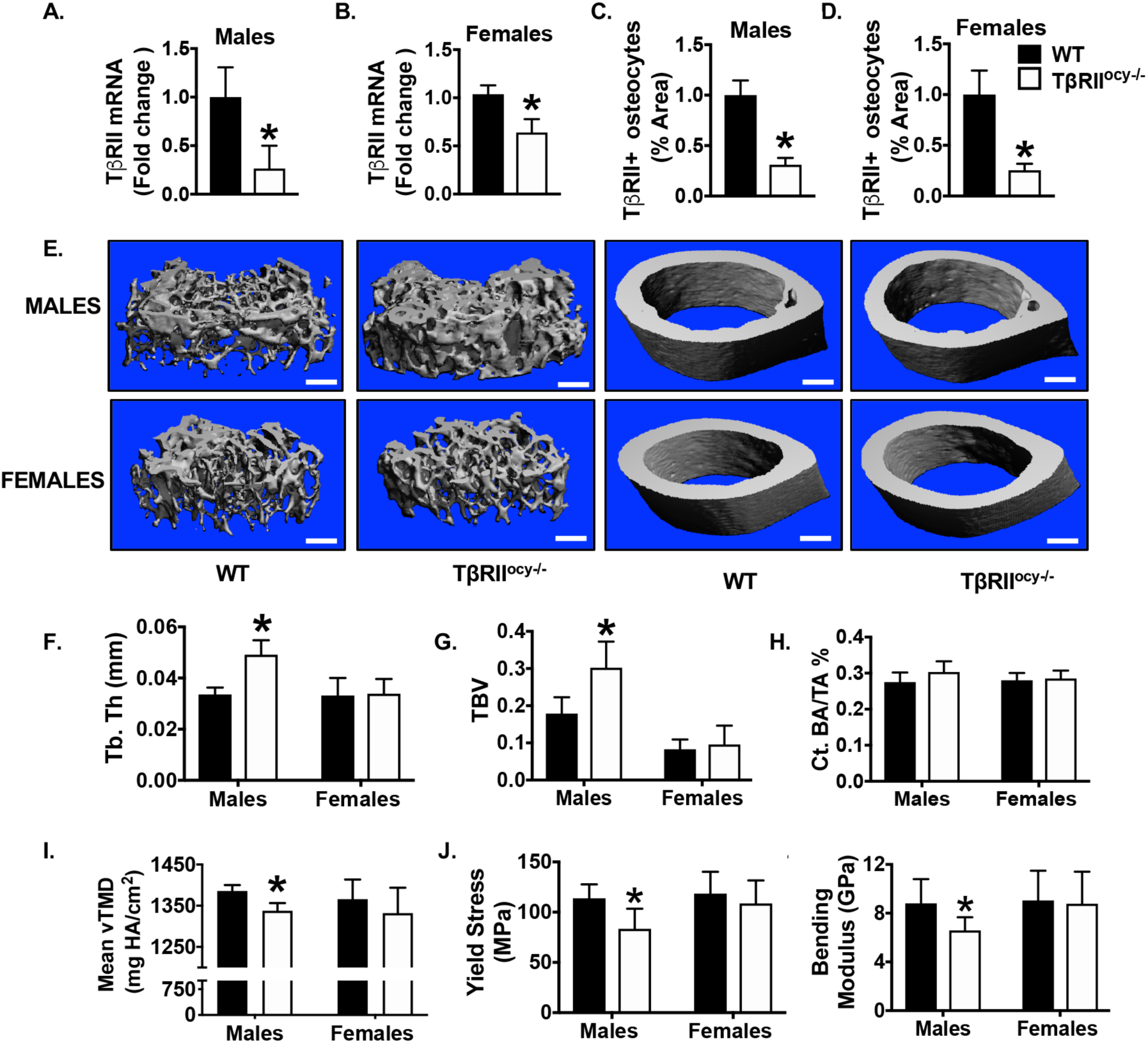
Loss of osteocyte-intrinsic TGFβ signaling impacts bone mass and quality in male mice but does not affect female mice. In male and female TβRII^ocy−/−^ mice, knockdown of TβRII at mRNA (A, B) (n=6-9 mice/ group) and protein (C, D) (n=4-6 mice/ group) level was assessed. μCT analysis was conducted on femurs from WT and TβRII^ocy−/−^ male and female mice (15-week-old). Representative μCT reconstructions of trabecular (left) and cortical (right) bone (E) from mice are shown. Trabecular bone thickness (Tb. Th.) (F) and bone volume fraction (BV/TV) (G), and cortical bone area (BA/TA) (H) and mineralization (Ct. Min) (I) were assessed (n=7-8 mice/group). Flexural testing of femurs from male and female WT and TβRII^ocy−/−^ mice shows bone quality parameters yield stress (J) and bending modulus (K) (n = 7 mice/group). Data for A-D are presented as mean SEM and as mean SD for E-K. *p < 0.05 different from WT group from Student’s t test. Scale bar for E is 100 μm (n = 7–8 mice/group).

**Figure 2.**
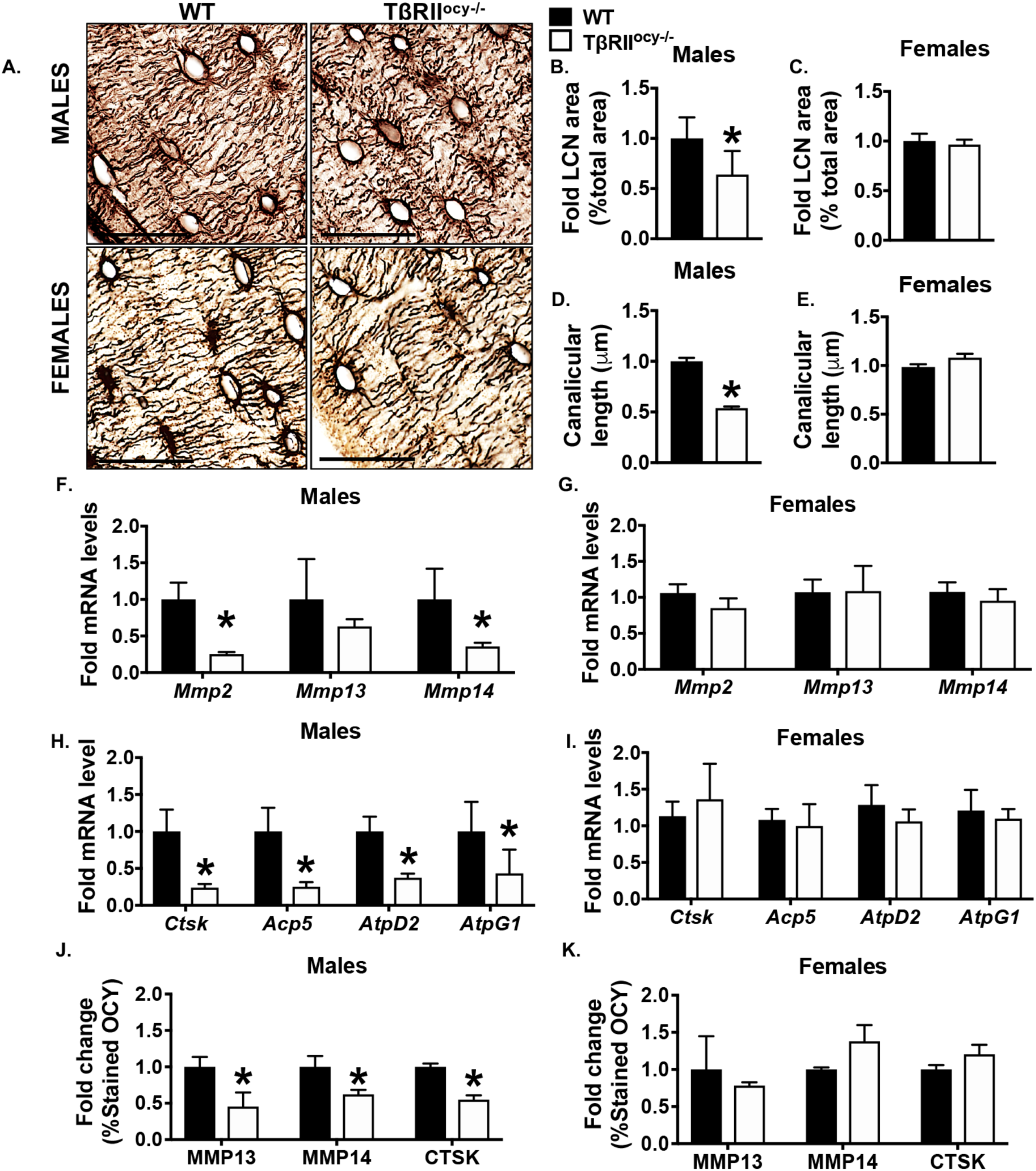
Osteocyte-specific disruption of TβRII reduces PLR genes in male but not female mice. Bones from 15-week-old male and female WT and TβRII^ocy−/−^ mice were harvested and processed for mRNA and protein analysis. Osteocyte lacuno-canalicular network (LCN) in the femoral cortical bones was assessed by silver nitrate staining. Representative images (A) and quantification of LCN area (B, C) and canalicular length (D, E) are shown. Scale bar, 20 μm; n=4-6 mice/group and 4 ROI/mouse. Expression of PLR genes Mmp2, Mmp13 and Mmp14 (F, G) and Ctsk, Acp5, Atp6v0d2 (AtpD2) and Atp6v1g1 (AtpG1) are shown (H-I) (n=6-9 mice/ group). Immunohistochemistry (IHC) for MMP13, MMP14, and CTSK was conducted on femoral cortical bones of male and female WT and TβRII^ocy−/−^ mice and percent positively stained osteocytes was quantified (J-K) (n = 3-4 mice/group and 4 ROI/mouse). Scale bar, 30 μm. Error bars indicate mean ± SEM, *p < 0.05 different from WT group from Student’s t test.

**Figure 3.**
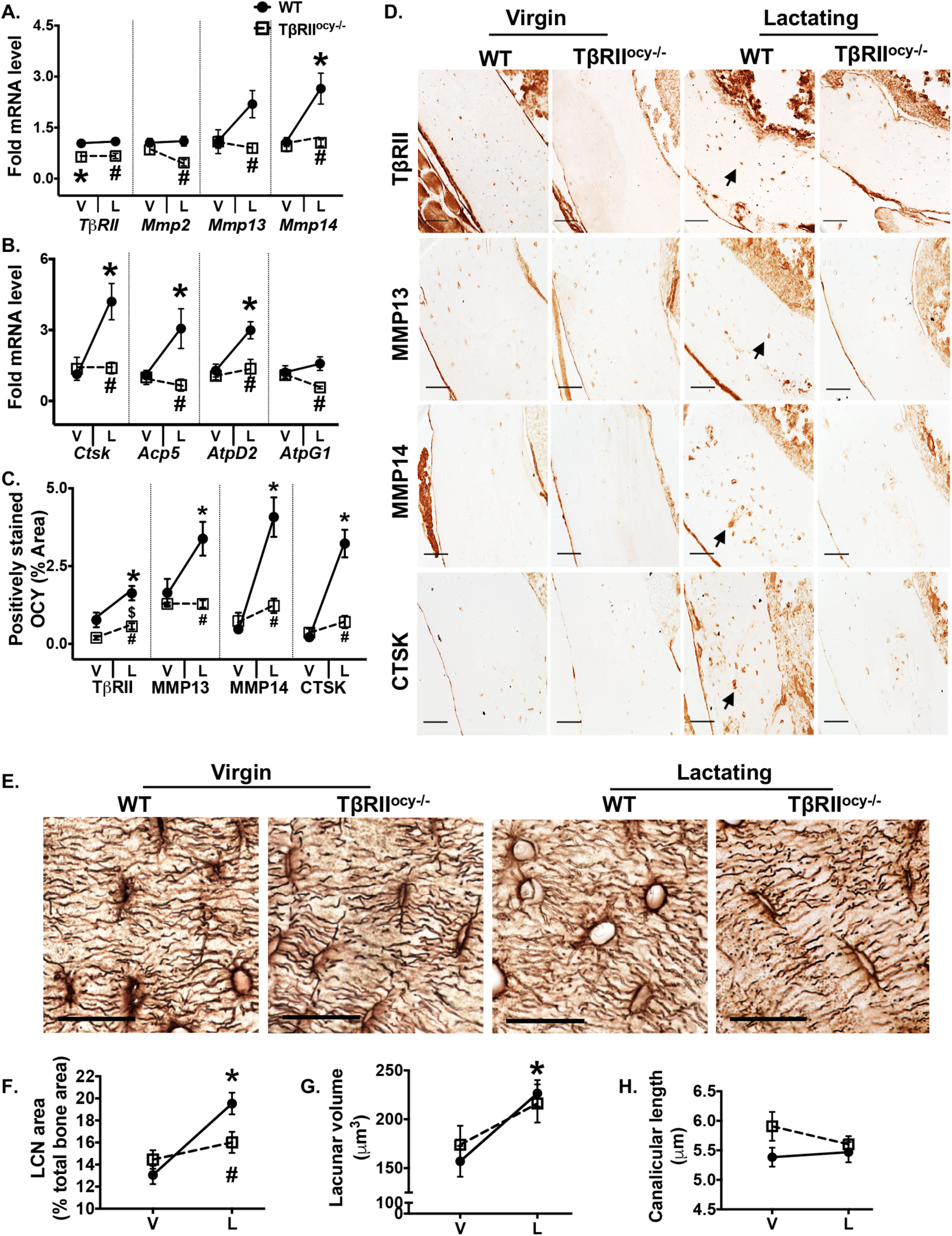
Lactation induced PLR blocked by osteocyte-specific disruption of TβRII gene. Bones from 15-week-old virgin and lactating WT and TβRII^ocy−/−^ female mice were harvested and processed for gene expression (A-B) and immunohistochemistry (C-D). qPCR analysis of TβRII and PLR genes Mmp2, Mmp13, Mmp14, Acp5, Ctsk, Atp6v0d2 and Atp6v1g1 (A-B) in virgin and lactating WT and TβRII^ocy−/−^ mouse bones was compared (n=6-9 mice/ group). Immunohistochemistry (IHC) for MMP13, MMP14, and CTSK was conducted on femoral cortical bones of virgin and lactating WT and TβRII^ocy−/−^ mice. Arrows in the representative images indicate positively stained osteocytes (D) that were quantified and normalized to total bone area (C) (n = 3-4 mice/group and 4 ROI/mouse). Change in osteocyte lacunar area (E, F) and canalicular length (H) of virgin and lactating WT and TβRII^ocy−/−^ mice were measured from Ploton silver stained cortical bone. Lacunar volume (G) was examined by SRμT (n=4-5 mice/group). Scale bar for D is 50 μm and 20 μm for E. Error bars indicate mean ± SEM, *p<0.05 compared to WT virgins, #p<0.05 compared to WT lactation group, and $ p<0.05 compared to TβRII^ocy−/−^ virgin, as calculated from one-way ANOVA with Tukey post-hoc test.

To assess the skeletal phenotype of TβRII^ocy−/−^ male and female mice, micro-computed tomography (μCT) was used. These analyses revealed an increase in trabecular thickness and trabecular bone volume in male TβRII^ocy−/−^ mice compared to the WT littermates (Figure 1E-G, table 1). Although cortical bone mass was unchanged, the cortical bone mineralization of TβRII^ocy−/−^ male mice was reduced by 3.5% relative to WT controls (Figure 1E, H-I). Flexural strength tests showed that both yield stress and bending modulus of TβRII^ocy−/−^ male bones was decreased by 20%, thereby indicating a reduction in the bone material quality of TβRII^ocy−/−^ males compared to WT mice (Figure 1J-K). As expected, based on our prior work on 8-week old TβRII^ocy−/−^ males, the current findings confirm the critical role of osteocyte-intrinsic TGFβ signaling in maintaining bone quality in 15-week old male mice.

None of these bone quality defects in TβRII^ocy−/−^ males were present in female TβRII^ocy−/−^ mice. In particular, the trabecular and cortical bone mass and geometry of TβRII^ocy−/−^ females remained intact, and the flexural strength of TβRII^ocy−/−^ female bones was identical to that of the WT group (Figure 1E-I, J-K, Table 2). These observations suggest that female mouse bones do not depend on osteocytic TGFβ signaling for maintaining bone mass and quality, and that osteocyte-intrinsic TGFβ impacts bone in a sexually dimorphic manner.

**Table 2:**
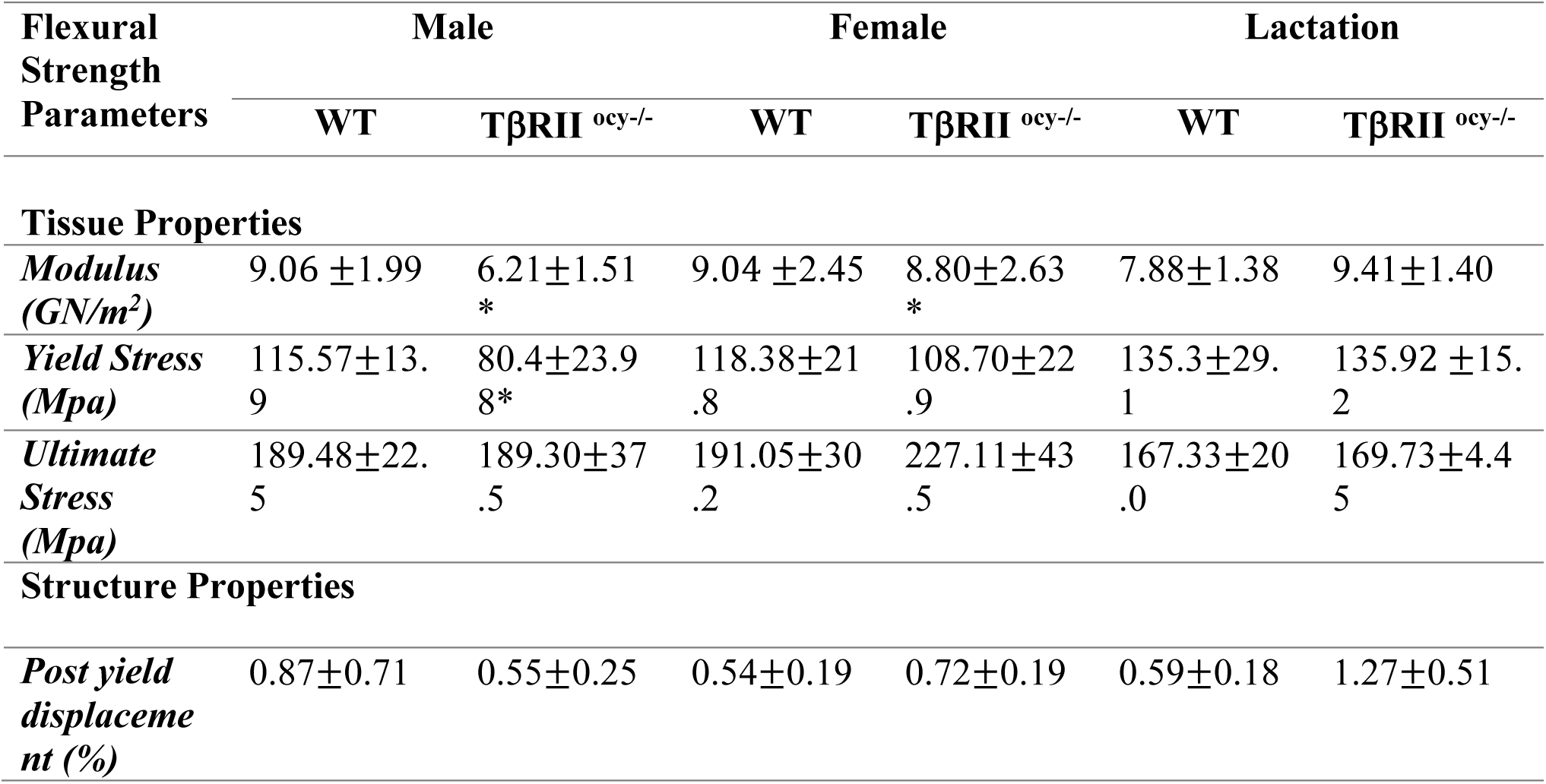
Flexural strength test derived macro-mechanical properties of 15 weeks-old male and virgin and lactating female WT and TβRII^ocy−/−^ mice. *p<0.05 significantly different from the male WT virgin group (Student’s t-test)

### 4.2 TGFβ signaling regulates osteocyte-mediated PLR in sex-specific manner

Bone quality is in part regulated by PLR, and our prior evidence from 8-week-old TβRII^ocy−/−^ male mice ascribes their defects in bone quality to PLR suppression^(32)^. Therefore, we hypothesized that the distinct bone quality phenotype of male and female TβRII^ocy−/−^ mice is driven by differences in osteocytic PLR. Since an intact lacuno-canalicular network (LCN) is a hallmark of functional PLR, we first examined the LCN in femora from 15-week old male and female TβRII^ocy−/−^ mice. Compared to WT males, the LCN of 15-week old TβRII^ocy−/−^ male bone was severely deteriorated, with a 50% reduction in LCN area and blunted canalicular length (Figure 2A-B, D). In addition, the male TβRII^ocy−/−^ bones also displayed a strong coordinated decrease in the expression of genes encoding factors involved in bone resorption during PLR, namely, Mmp2, Mmp13, Mmp14 (matrix metalloproteinases), Ctsk (Cathepsin K), Acp5 (tartrate-resistant acid phosphatase), Atp6v0d2 (AtpD2), and Atp6v1g1 (AtpG1) (vacuolar ATPase pump encoding genes) (Figure 2F and H). Moreover, osteocyte-intrinsic protein expression of MMP13, MMP14 and CTSK was reduced in TβRII^ocy−/−^ male mouse bones (Figure 2J).

Female TβRII^ocy−/−^ mice responded differently to osteocyte-intrinsic TGFβ ablation. In the female TβRII^ocy−/−^mice, the osteocyte LCN was intact with no changes observed in LCN area or canalicular length (Figure 2A, C and E). Consistent with these results, the expression of PLR genes in the bone of female TβRII^ocy−/−^ mice did not differ from that in WT controls (Figure 2G, I and K). Thus, the homeostatic regulation of PLR and bone quality in female mice occurs independently of TGFβ signaling. Since the absence of osteocytic TGFβ signaling in males adversely affects these outcomes, our data show a sexually dimorphic role for TGFβ in the control of PLR and bone quality.

### 4.3 Osteocytic deletion of TβRII prevents lactation induced PLR

Although the homeostatic regulation of PLR in females is TGFβ-independent, we hypothesized that intact TGFβ signaling may be required to induce PLR in response to the metabolic stress of lactation. Using an established murine model of lactation, we assessed PLR outcomes in 15-week old lactating TβRII^ocy−/−^ and WT mice. These results were compared to those in the 15-week old female TβRII^ocy−/−^ and WT virgin controls, described above. Upon validation of targeted TβRII deletion at the mRNA and protein levels in the lactating TβRII^ocy−/−^ mice (Figure 3A, C and D), we monitored the effect of lactation on the expression of PLR genes. WT mice showed a 2.5 to 4-fold induction in the expression of the Mmp14, Ctsk, Acp5 and Atp6v0d2 genes (Figure 3A-B) during lactation. The increased expression of PLR enzymes was also apparent at the protein level, with a 3 to 5-fold increase in the percentage of MMP13, MMP14 and CTSK-positive osteocytes in the lactating WT group (Figure 3C-D). Consistent with these observations, WT lactating mice undergo a 50% increase in the LCN area and a corresponding 40% increase in the mean lacunar volume relative to WT virgin controls (Figure 3E-H).

On the contrary, lactation failed to induce PLR in TβRII^ocy−/−^ mice. In particular, lactating TβRII^ocy−/−^ mice did not exhibit a significant induction in PLR genes, LCN area, lacunar volume and canalicular length (Figure 3 A-H). Similar to resorptive genes, the lactation-induced expression of bone formation genes, Dmp1 and Runx2, was absent in TβRII^ocy−/−^ mice (Figure S2A-B). Therefore, distinct from the TGFβ-independent regulation of PLR in female bone at homeostasis, osteocyte-intrinsic TGFβ signaling is required for the induction of PLR during lactation.

### 4.4 Targeted deletion of osteocytic TβRII mitigates lactation-induced bone loss

µCT analyses revealed that during lactation, WT mice lose more than 52% of trabecular bone volume fraction (BV/TV, Table 1). This decrease in BV/TV is reflected by the corresponding reduction in trabecular number (28%) and thickness (32%) and increase in trabecular spacing (45%) during lactation in WT mice (Table 1, Figure 4A-B). The trabecular bone mineralization is also reduced by 9% with lactation in WT mice. A decrease in cortical bone thickness (27% reduction) and mass (25% reduction) was observed in lactating WT mice, but the cortical mineralization remained intact (Table 1, Figure 4A-C). In contrast to WT mice, TβRII^ocy−/−^ mice did not lose trabecular bone mass during lactation, and their trabecular number and spacing remained unaltered. TβRII^ocy−/−^ mice lost some cortical thickness and bone mass during lactation, however the magnitude of change was significantly lower in lactating TβRII^ocy−/−^ mice than WT mice (i.e. 14% vs. 25%, p<0.05) (Figure 4A-C).

**Figure 4.**
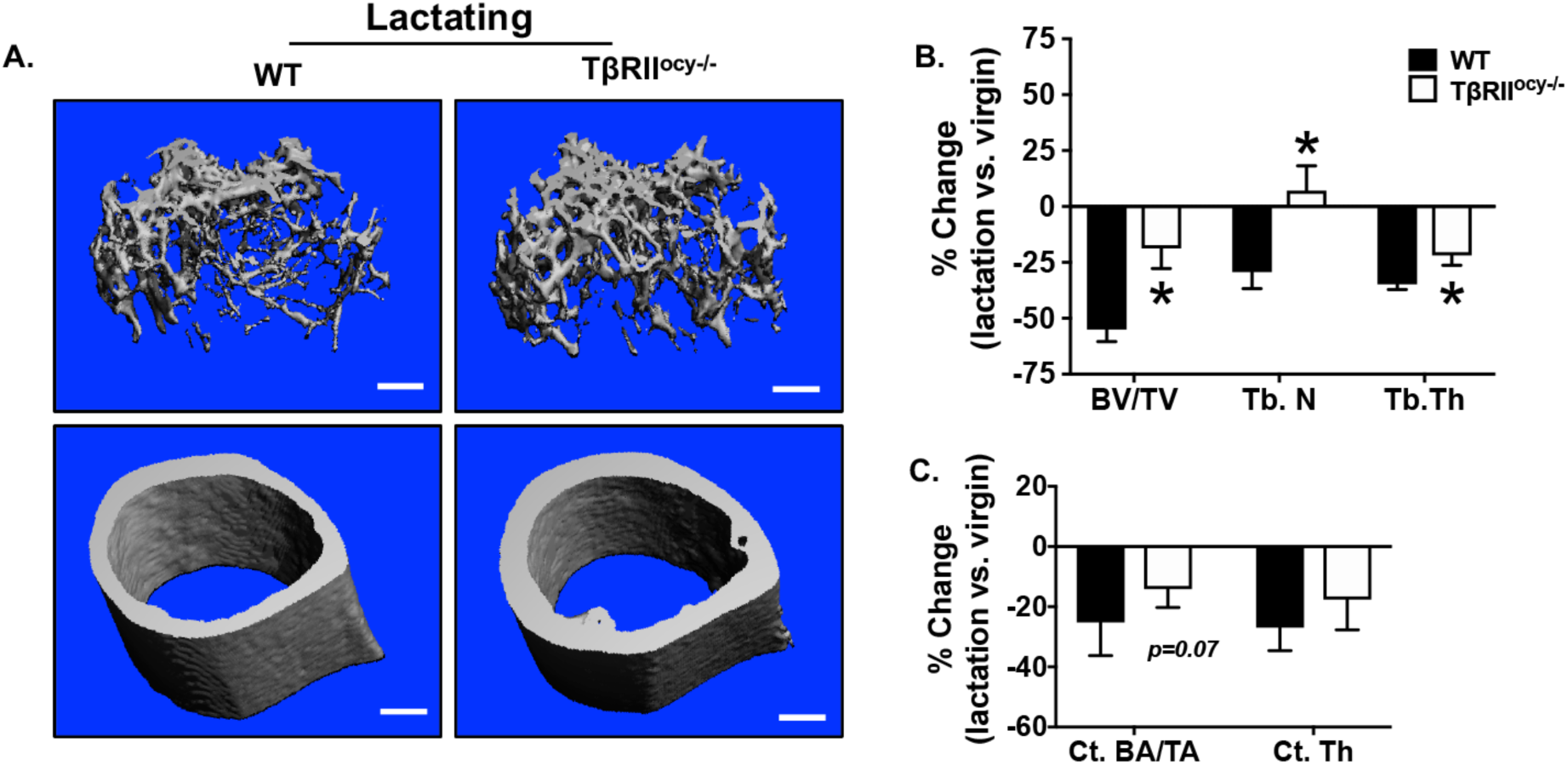
Lactation induced trabecular bone mass in lactating TβRII^ocy−/−^ is mitigated. μCT analysis of femurs from 15-week old WT and TβRII^ocy−/−^ lactating female mice. Representative μCT reconstructions show trabecular (top) and cortical (bottom) bones (A). Decline in trabecular bone parameters: trabecular thickness (Tb. Th.), mineralization (Tb. BMD) and bone volume fraction (BV/TV) (B), and cortical bone parameters: cortical thickness (Ct. Th), and cortical bone volume (C) with lactation is represented as percent change compared to the respective virgin groups for both genotypes. Scale bar, 100 μm (n = 6-10 mice/group). Error bars indicate mean ± SD, *p<0.05 compared to WT lactation as calculated from Student’s t-test.

The high-resolution quantitative imaging available through synchrotron SRµCT further revealed a comparable trend towards increased mineralization around osteocyte lacunae with lactation that was apparent in both genotypes (Figure S3). However, µCT data did not show significant differences in overall cortical bone mineralization during lactation in either genotype (Table 1). Accordingly, bone bending modulus, yield stress, and ultimate stress were unaffected by lactation in either genotype (Table 2). Collectively, these data establish the role of osteocyte-intrinsic TGFβ signaling in lactation, such that absence of this mechanism alleviates lactation-induced bone remodeling by osteocytes and surface cells.

### 4.5 PTH involvement in the sexually dimorphic effects of osteocytic TGF β

As we previously reported, the elevated bone mass in 8-week old TβRII^ocy−/−^ male mice is accompanied by reduced Rankl (Tnfsf11) expression and decreased osteoclast numbers^(32)^. Therefore, we hypothesized that the sex-specific differences in bone mass in TβRII^ocy−/−^ male and female mice were driven by dimorphic regulation of Rankl. Here we find that 15-week old male TβRII^ocy−/−^ mice also express low levels of Rankl mRNA, with a stark reduction in the Rankl/Opg ratio, compared to WT males. Female TβRII^ocy−/−^ mice, on the other hand, showed intact Rankl and Rankl/Opg ratios during homeostasis (Figure 5A-B). During lactation, the high demand for calcium drives bone resorption, in part, through robust Rankl expression^(43)^. Indeed, lactating WT mice increase Rankl mRNA expression with a resulting 7-fold increase in the Rankl/Opg ratio (Figure 5C and S2C). Lactating TβRII^ocy−/−^ mice, however, fail to stimulate Rankl mRNA expression. Instead Opg mRNA levels are suppressed in the lactating TβRII^ocy−/−^ mice, through an apparent compensatory mechanism to increase the Rankl/Opg ratio 2-fold to release calcium for lactation (Figure 5C and S2C).

**Figure 5.**
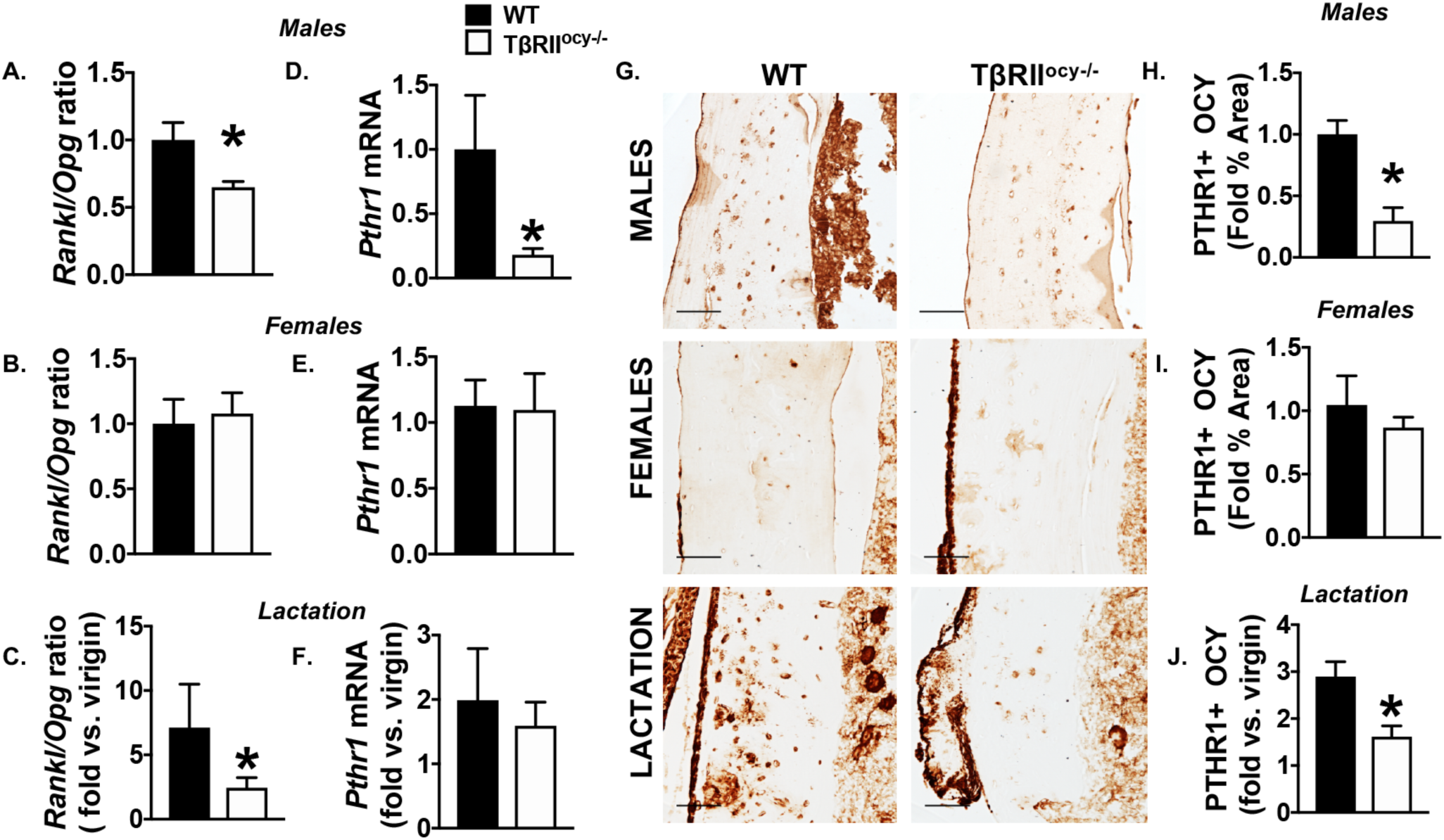
PTHR1 dependent signaling is reduced in TβRII^ocy−/−^ males and lactating females. qPCR analysis for Rankl and Opg in WT and TβRII^ocy−/−^ males (A) and virgin (B) and lactating (C) females is presented as a ratio of Rankl/Opg mRNA, with lactating groups shown relative to corresponding virgin groups. Pthr1 mRNA was also quantified by qPCR(D-F) and by immunohistochemistry (G-J). Percentage of osteocytes that stained positive for PTHR1 in bones of WT and TβRII^ocy−/−^ male and virgin and lactating female mice were quantified and normalized against total bone area as shown (H-J) (n=3-4 mice/group, 4 ROI/mouse). Error bars indicate mean ± SEM, *p<0.05 compared to respective WT group as calculated from Student’s t-test.

These differences in Rankl/Opg expression led us to focus on the role of PTH signaling in the sexually dimorphic effects of TGFβ in osteocytes. Several lines of evidence motivated our focus: First, PTH signaling directly regulates Rankl gene expression in osteocytes^(44,45)^. Second, PTH and parathyroid hormone related peptide (PTHrP), key regulators of calcium and phosphate homeostasis, induce PLR during lactation^(5,21,46–48)^. Third, PTH and PTHrP signal through a common parathyroid hormone receptor (PTHR1) in osteocytes; and PTHR1 ablation in osteocytes blocks lactation-inducible PLR^(21)^. Fourth, as in PTHR1^ocy−/−^ mice, TβRII^ocy−/−^ mice are unable to induce PLR or bone loss during lactation^(21)^. Fifth, TβRII and PTHR1 interact in osteoblasts to augment PTH signaling and increase bone formation^(49)^. Based on these findings, we hypothesized that TGFβ and PTH signaling utilize a common molecular mechanism to regulate PLR.

To test this hypothesis, we first examined the effect of TβRII ablation on the expression of PTHR1 in the osteocytes. Both immunohistochemistry and quantitative PCR revealed a stark reduction (>70%) in the expression of PTHR1 in osteocytes of TβRII^ocy−/−^ male mice compared to WT (Figure 5D, G, H). This was consistent with the low Rankl mRNA and Rankl/ Opg ratios seen in the TβRII^ocy−/−^ males. In females during homeostasis, no changes were detected in the expression of PTHR1 at the mRNA or protein level between WT and TβRII^ocy−/−^ mice, which agrees with the lack of changes in Rankl/Opg in TβRII^ocy−/−^ females (Figure 5E, G, I). During lactation, PTHR1 expression is robustly increased in WT females, but loss of osteocytic TβRII mitigates this response in lactating TβRII^ocy−/−^ mice (Figure 5F, G, J), consistent with the lower overall Rankl/Opg ratio in lactating TβRII^ocy−/−^ mice compared to lactating WT mice.

Lastly, we evaluated serum levels of calcium, phosphate, and PTH in each group (Table 3). Phosphate levels did not differ in any condition. The two groups that had reduced PTHR1 expression, male and lactating female TβRII^ocy−/−^ mice, had elevated levels of serum calcium relative to WT controls. In male TβRII^ocy−/−^ mice, the hypercalcemia may result from increased serum PTH levels. Importantly, in non-lactating female mice, none of these parameters differed between TβRII^ocy−/−^ mice and WT controls, which is consistent with the insensitivity of these mice to osteocyte-intrinsic TGFβ ablation in all the other outcomes we analyzed. Taken together these findings suggest a key role of TβRII in regulating PTHR1 expression in osteocytes, and that this coordination between TGFβ and PTH signaling dictates the PLR response in osteocytes during homeostasis and metabolic stress.

**Table 3:** Serum analysis of 15 weeks-old male and female WT and TβRII^ocy−/−^ mice. *p<0.05 significantly different from the male WT virgin group (Student’s t-test) ^a^ p<0.05 significantly different from the WT virgin group ^b^ p<0.05 significantly different from the KO virgin group

| Serum | Male |  | Female |  | Lactation |  |
| --- | --- | --- | --- | --- | --- | --- |
|  | WT | TβRII<sup>ocy-/-</sup> | WT | TβRII<sup>ocy-/-</sup> | WT | TβRII<sup>ocy-/-</sup> |
| **PTH (pg/ml)** | 269.17±59.59 | 484.5±71.7<br>1* | 225.01±48.93 | 222.63±19.62 | 41.88±5.09<sup>a</sup> | 39.1±8.70<sup>a,b</sup> |
| **PTHrP (pg/ml)** | 293.15±55.9 | 283.58±62.21 | 346.50±36.72 | 311.78±39.31 | 183.24±55.3<sup>a</sup> | 222.37±33.5<sup>a,b</sup> |
| **Calcium (mg/dl)** | 6.84±0.83 | 9.49±1.55* | 5.22±1.33 | 5.54±0.74 | 3.62±2.63<sup>a</sup> | 7.86±1.5<sup>a,b</sup> |
| **Phosphate (mg/dl)** | 20.21±1.58 | 19.13±1.60 | 14.26±1.58 | 12.89±1.49<br>5 | 13.18±4.33 | 14.61±6.5 |

## 5. Discussion

This study provides insight into sex-specific variation in bone fragility by revealing dimorphism in the regulation of osteocyte perilacunar/ canalicular remodeling (PLR). Using a genetic model of osteocyte-specific deletion of TGFβ receptor II (TβRII), we show that loss of osteocyte-intrinsic TGFβ signaling affects the skeleton of male, but not female, mice in homeostatic conditions. Male TβRII^ocy−/−^ mice have defective bone quality, despite normal bone mass, resulting from impaired osteocytic PLR with deteriorated canalicular networks and reduced expression of requisite PLR enzymes. On the contrary, female TβRII^ocy−/−^ mice are protected from these deficits in both PLR and bone quality. Osteocytic TGFβ signaling is necessary for PLR induction in female mice during lactation. Lactation-induced PLR is mitigated in TβRII^ocy−/−^ females, as is lactation-induced bone loss. These findings highlight distinct roles for TGFβ signaling in regulating PLR in males and females to maintain bone homeostasis and implicate TGFβ as an essential regulator of PLR in females principally during the metabolic stress of lactation.

Gender disparity in fracture risk stems from differences in peak bone mass accrual, bone size, and geometry^(50)^. Sex steroids that essentially drive skeletal variations in men and women integrate systemic cues such as that from vitamin D, IGF/GH, and PTH to dictate dimorphism at the cellular level^(19,51–53)^. Indeed, sex differences in the cellular function of osteoblasts and osteoclasts have been attributed to differences in the bone phenotype of male and female mice^(17–20)^. Differences in bone quality also contribute to the gender disparities in fracture risk. Osteocyte PLR is one cellular mechanism that plays a pivotal role in maintaining bone quality. Although the contribution of PLR to sex differences in bone quality was unknown, osteocytes clearly have a dimorphic response to aging^(54,55)^. The osteocyte lacuno-canalicular network, which is maintained by PLR, deteriorates more rapidly in aging females than in aging males^(54,55)^. Here we describe sexually dimorphic regulation of osteocytes in the control of PLR and bone quality by TGFβ signaling. Whereas in males, loss of osteocytic TGFβ signaling suppresses PLR and increases bone fragility, female TβRII^ocy−/−^ mice demonstrate a complete lack of this regulatory mechanism. Such skeletal adaptations in females could be one of the mechanisms by which the female skeleton compensates to meet the skeletal demands of pregnancy and lactation. The extent to which this ‘protective mechanism’ in reproductive age females is lost following menopause, or if it contributes to the increased risk of post-menopausal bone fragility remains unknown. Nevertheless, our study shows that sex specific differences in osteocyte sensitivity to TGFβ signaling contribute to the sexual dimorphism in PLR and bone quality.

TGFβ signaling has previously been implicated as one of the mechanisms recruited to confer sex specific differences in bone. An intricate crosstalk exists between TGFβ signaling and estrogen in bone. Estrogen promotes de novo synthesis and activation of TGFβ ligand in bone. Conversely, the loss of estrogen in ovariectomized rats leads to reduced TGFβ expression in bone^(56–60)^. TGFβ, in turn, mediates bone protective effects of estrogen by increasing osteoclast apoptosis^(57,61)^. With systemic inhibition of TGFβ signaling using a pharmacologic TGFβ receptor I kinase inhibitor (TβRI-I), we observed increased sensitivity of the female skeleton to TGFβ inhibition, with increased osteoblast proliferation and bone mass accrual in female mice than males^(36)^. Here, in osteocytes of female mice with ablated TβRII, we observed that loss of osteocytic TGFβ signaling has no impact on PLR gene expression, the osteocyte lacuno-canalicular network, and bone quality. The extent to which these sex specific differences in osteocyte TGFβ signaling are related to estrogen levels remains to be determined. The remarkably distinct regulation of TGFβ signaling by sex-specific factors in osteoblasts, osteoclasts and osteocytes could afford independent, yet coordinated, regulation of bone quantity and quality to maintain mineral homeostasis and bone strength during hormonal and reproductive cycles in females.

Systemic calcium homeostasis is tightly regulated by the parathyroid gland that controls the interplay between the calciotropic hormones like, parathyroid hormone (PTH), parathyroid hormone related protein (PTHrP), calcitonin, and calciferol (1α,25-dihydroxycholecalciferol, vitamin D3). Together these hormones establish an equilibrium between dietary calcium absorption through the intestines, calcium reabsorption and excretion through kidneys, and its storage and release from bones^(62)^. In bone, osteocytes show particularly high sensitivity to calcium fluctuations and are a target of PTH action^(21,63,64)^. PTHR1, the receptor that binds both PTH and PTHrP, is highly expressed in bone and kidney and mediates the PTH-dependent regulation of calcium homeostasis. Although, little is known about the crosstalk of PTH and PTHrP with TGFβ signaling in bone, PTHR1 interacts directly with TβRII. In osteoblasts, PTHR1 and TβRII coordinately modulate bone formation^(49)^. In osteocytes, however, we observed that ablation of TβRII causes a marked reduction in PTHR1 expression and Rankl/ Opg ratios only in male mice. Given that PTHR1 is a key regulator of PLR, the PLR and bone quality defects observed in the male TβRII^ocy−/−^ may result from reduced PTHR1-dependent signaling. The reduced PTHR1-dependent signaling in TβRII^ocy−/−^ male mice appears to be compensated by increased systemic PTH levels, which in turn increase systemic calcium levels, likely by targeting tissues other than bone.

In female mice, the role of TGFβ in osteocytes differs dramatically between homeostatic conditions and the metabolic stress of lactation. In homeostatic conditions, female TβRII^ocy−/−^ mice differ from TβRII^ocy−/−^ males, in that they show intact osteocytic PTHR1 expression and PLR. Other pathways that are active in females, but not in males, may converge on PTHR1 to sustain its expression in female mice at homeostasis. This capacity of female TβRII^ocy−/−^ osteocytes to normalize PTHR1 levels at homeostasis may contribute to the intact bone quality of female TβRII^ocy−/−^ mice. This capacity is challenged by the metabolic stress of lactation, which normally induces PLR to meet the high calcium demands for breastmilk production. PLR induction during lactation occurs in a hormonal milieu with low PTH and estradiol, but with high PTHrP levels. While we demonstrated reduction in circulating PTH with lactation in both genotypes, the circulating level of PTHrP was not significantly increased in either genotypes, most likely due to insufficient sensitivity of our assay. PLR induction during lactation also requires osteocytic PTHR1. Although PTHR1 mRNA is induced in lactation in both genotypes, the induction of PTHR1 protein in osteocytes is abrogated in lactating TβRII^ocy−/−^ mice relative to WT littermates. Accordingly, with reduced osteocytic PTHR1 protein levels, PLR induction in female TβRII^ocy−/−^ mice during lactation is blocked. Our results point to the possibility that the level of PTHR1 is a pivotal determinant of the effect of TGFβ on PLR. Thus, the differential PLR response of osteocytes in the TβRII^ocy−/−^ mice in males vs. females, and in homeostasis vs. metabolic stress, may result from differences in the ability to stimulate PTHR1-mediated bone resorption by osteocytes. It is likely that such molecular mechanisms are in place to compensate for reproductive bone loss and protect the female skeleton from higher fracture risk. Another such compensatory mechanism that was recently suggested includes accrual of higher trabecular bone mass than mechanically necessary by female rats compared to males^(1,65)^.

In conclusion, our study highlights sexual dimorphism in TGFβ’s regulation of osteocyte perilacunar/canalicular remodeling activity. With implication of PLR in both physiological and pathological conditions, including renal osteodystrophy, glucocorticoid-induced osteonecrosis, osteoarthritis, and hyperparathyroidism^(21,66–71)^, defining the sex differences in these cell-intrinsic cues and molecular networks in osteocytes will be critical to address sexual dimorphism in the prevalence and manifestation of bone fragility, as well as refining therapeutics for reducing fracture risk.

## 6. Author Contributions

Study design, N.S.D and T.A.; Study conducted, N.S.D., C.S.Y., C.M.M., and C.A.; Data collection and Analysis, all authors; Data interpretation: N.S.D and T.A.; Writing – Original Draft, N.S.D.; Revising manuscript and approving final version of manuscript: all authors; Supervision, T.A.; Project Leadership, N.S.D. and T.A.; Funding Acquisition, T.A. N.S.D and T.A. take responsibility for the integrity of the data analysis.

## 7. Acknowledgments

The authors gratefully acknowledge J.J. Woo for expert technical assistance. This research was supported by NIH-NIDCR grant R01 DE019284 (T.A.), including a supplement from the Office of Research on Women’s Health, Department of Defense (DoD) grant PRORP OR130191 (T.A.), NSF grant 1636331, NIH-NIAMS grant R21 AR067439, NIH-NIAMS grant P30 AR066262-01 (T.A.), and the Read Research Foundation (T.A.). The authors acknowledge the use of the x-ray synchrotron beamlines 8.3.2 at the Advanced Light Source (ALS) at LBNL. The ALS is supported by the Director (Office of Science, Office of Basic Energy Sciences) of the U.S. Department of Energy under contract DE-AC02-05CH11231. We are grateful for helpful conversations with our colleagues in the San Francisco Veteran’s Administration Medical Center (SF-VAMC) Endocrine Unit throughout the development of this project.

