## Supplemental Data for "TGFβ regulation of perilacunar/canalicular remodeling is sexually dimorphic"

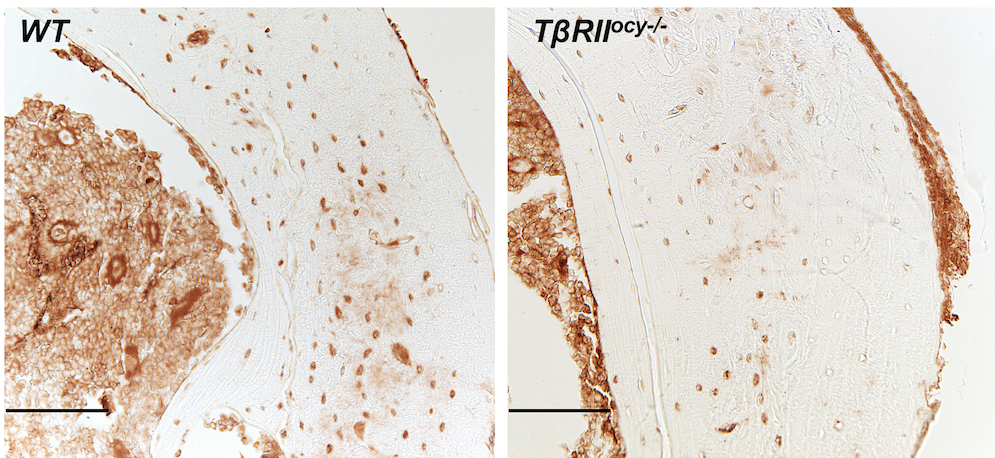

**Figure S1:** Representative image of TβRII-stained osteocytes (scale bar, 100 μm) in the femoral cortical bone from WT and TβRII^ocy−/−^ mice (15-week-old males) (n = 4-6 mice/group).

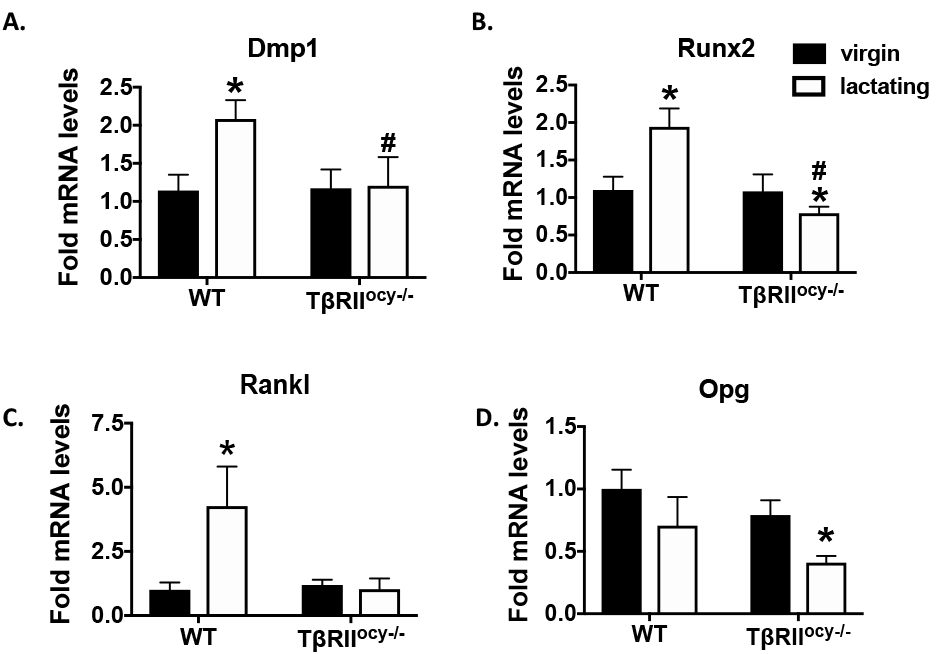

**Figure S2:** qPCR analysis of Dmp1 (A), Runx2 (B), Rankl (C), and Opg (D) in WT and TβRII^ocy−/−^  virgin and lactating female mice bones is shown (A-B). (n = 6-9 mice/ group).

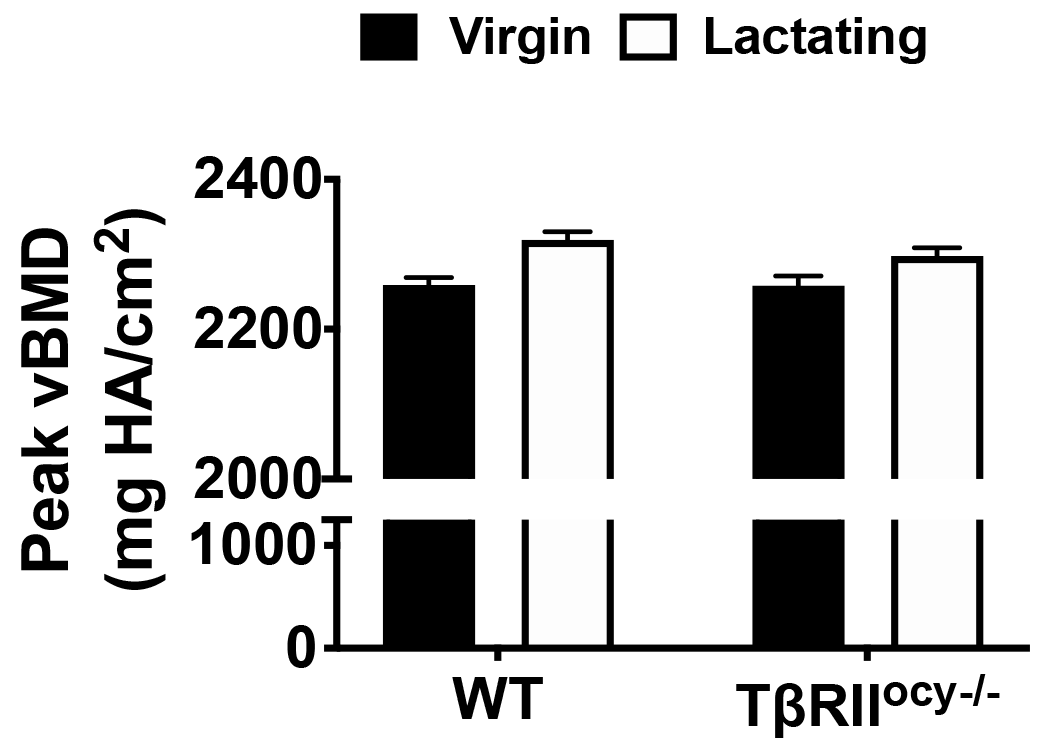

**Figure S3:** SRμT shows mineralization of osteocyte lacunae of WT and TβRII^ocy−/−^ bones harvested from virgin and lactating mice (n = 3–4 mice/group).

**Table S1: Primer sequences used for qPCR analysis.**

| **Gene** | **Sequence** |
| --- | --- |
| 18s RNA (sense)  18s RNA (antisense) | 5’-CGAACGTCTGCCCTATCAAC-3’  5’-GGCCTCGAAAGAGTCCTGTA-3’ |
| Mmp13 (sense)  Mmp13 (antisense) | 5’-CGGGAATCCTGAAGAAGTCTACA-3’  5’-CTAAGCCAAAGAAAGATTGCATTTC-3’ |
| Mmp14 (sense)  Mmp14 (antisense) | 5’-AGGAGACGGAGGTGATCATCATTG-3’  5’-GTCCCATGGCGTCTGAAGA-3’ |
| Mmp2 (sense)  Mmp2 (antisense) | 5’-AACGGTCGGGAATACAGCAG-3’  5’-GTAAACAAGGCTTCATGGGG-3’ |
| Ctsk (sense)  Ctsk (antisense) | 5’- GAGGGCCAACTCAAGAAGAA-3’  5’- GCCGTGGCGTTATACATACA-3’ |
| Acp5 (sense)  Acp5 (antisense) | 5’-CGTCTCTGCACAGATTGCAT-3’  5’-AAGCGCAAACGGTAGTAAGG-3’ |
| Rankl (sense)  Rankl (antisense) | 5’-CCAAGATCTCTAACATGACG-3’  5’-CACCATCAGCTGAAGATAGT-3’ |
| Opg (sense)  Opg (antisense) | 5’-AGAGCAAACCTTCCAGCTGC-3’  5’-CTGCTCTGTGGTGAGGTTCG-3’ |
| Serpine1 (sense) Serpine1 (antisense) | 5’-CAGATGACCACAGCGGGGAA-3’  5’-GGCATGAGCTGTGCCCTTCT-3’ |
| Atp6v0d2 (sense)  Atp6v0d2(antisense) | 5’-TCTTGAGTTTGAGGCCGACAG-3’  5’-GCAACCCCTCTGGATAGAGC-3’ |
| Atp6v1g1 (sense)  Atp6v1g1 (antisense) | 5’-CCGTTCTCTCAGCCCAAAGT-3’  5’-CTCCGGTTCTTTCGCTTGC-3’ |
